# Development of a quantitative metagenomic approach to establish quantitative limits and its application to viruses

**DOI:** 10.1101/2022.07.08.499345

**Authors:** Kathryn Langenfeld, Bridget Hegarty, Santiago Vidaurri, Emily Crossette, Melissa Duhaime, Krista Wigginton

## Abstract

Quantitative metagenomic methods are maturing but continue to lack clearly-defined analytical limits. Here, we developed a computational tool, QuantMeta, to determine the absolute abundance of targets in metagenomes spiked with synthetic DNA standards. The tool establishes (1) entropy-based detection thresholds to confidently determine the presence of targets, and (2) an approach to identify and correct read mapping or assembly errors and thus improve the quantification accuracy. Together this allows for an approach to confidently quantify absolute abundance of targets, be they microbial populations, genes, contigs, or metagenome assembled genomes (MAGs). We applied the approach to quantify single- and double-stranded DNA viruses in wastewater viral metagenomes, including pathogens and bacteriophages. Concentrations of total DNA viruses in wastewater influent and effluent were greater than 10^8^ copies/mL using QuantMeta. Human-associated DNA viruses were detected and quantifiable with QuantMeta thresholds, including polyomavirus, papillomavirus, and crAss-like phages, at concentrations similar to previous reports that utilized quantitative PCR-based assays. Our results highlight the higher detection thresholds of quantitative metagenomics (∼500 copies/μL) as compared to PCR-based quantification (∼10 copies/μL) despite a sequencing depth of 200 million reads per sample. The QuantMeta approach, applicable to both viral and cellular metagenomes, advances quantitative metagenomics by improving the accuracy of measured target absolute abundances.

## INTRODUCTION

Metagenomics has provided unprecedented insights into the diversity, structure, and function of microbial communities across various environments, from natural and engineered systems to the human body (1). However, metagenomic data is inherently relative, which complicates direct comparisons of population and gene abundances across samples—particularly in systems with fluctuating biomass concentrations, such as wastewater treatment plants, or when comparing healthy and disease states. Developing a quantitative metagenomics approach that yields accurate absolute abundances for numerous targets simultaneously would greatly expand the scope of conclusions that can be drawn from these studies.

One approach to making metagenomic data quantitative is to convert relative abundances to absolute abundances using total cell estimates derived from flow cytometry or concentrations of 16S rRNA or housekeeping genes measured by quantitative PCR (2-5). However, these methods rely on time- and resource-intensive ancillary measurements that can be prone to errors. Additionally, the latter approach is unsuitable for virus quantification, as viruses lack conserved genes and are challenging to quantify using cytometry. An alternative approach involves adding DNA standards at known concentrations to samples, enabling absolute abundance measurements in metagenomes (6-15). Here, target gene or genome counts in the metagenome are quantified by comparing the number of reads mapping to these standards against the known standard concentrations (6,8,11-13,15).

Various spike-in standards can aid metagenomic quantification, including foreign genomes (12,13), mock microbial communities (15), and synthetic DNA sequences (6,8,11). Synthetic DNA standards offer unique advantages that make them increasingly popular: they avoid non-specific mapping with unique nonsense sequences (6,8), can be tailored to match expected microbiome characteristics (e.g., DNA lengths, GC content, and concentrations), and are readily available in premade mixtures (16). These standards have been used to examine temporal and spatial microbial variability in healthcare environments (9), track antibiotic resistance genes and pathogens in wastewater (10) and manure (11), and and analyze microbial community structure in saltmarshes (7). Additionally, they serve as positive controls in metagenomics, helping to establish false positive thresholds (17). As with any quantitative method, the next step is to define limits of detection and quantification to meet quality targets for bias, precision, and total error (18,19).

Despite the growing application of spike-in standards for quantification of targets in metagenomes, the approach lacks well-defined and standardized limits of detection and quantification (20), as is typical of other analytical methods (18). Ideally, detection limits would be established dynamically for a given dataset and target. Beyond detection, accurate target quantification relies on accurate read mapping to sequences of interest. Previous studies that implemented quantitative metagenomics determined target concentrations by mapping reads onto targets from databases (i.e., reference-based quantification) (7,9-11). Targets can also be quantified in metagenomes by mapping reads onto contigs assembled from the same sample (i.e., contig-based quantification). Ultimately, the average number of reads mapped to a sequence (read depth) is used to quantify a target; consequently, read mapping errors caused by non-specific mapping (i.e., reads mapping to a target other than its true origin (21)) or assembly errors (22) may reduce quantification accuracy. An approach to detect and, when possible, correct read mapping errors is needed to improve the accuracy of quantitative metagenomics.

In this study, we developed a computational tool, QuantMeta, to determine the absolute abundance of targets in metagenomes that have been spiked with synthetic DNA standards. Similar to the two-step process in quantitative PCR, where researchers first establish limits of detection and quantification before reliably applying them in a new system, our approach uses an analogous process to establish limits of detection and whether a target is quantifiable in metagenomics. QuantMeta incorporates (1) detection thresholds (acting similarly to limit of detection) to determine presence or absence of targets, (2) identification and correction of read mapping errors that affect accuracy of quantification, and (3) determination of absolute abundance of targets (Figure 1). We demonstrated the application of QuantMeta by quantifying ssDNA and dsDNA viruses in municipal wastewater samples. We anticipate that this approach will be useful in future quantitative metagenomics studies for both viral (virome) and entire microbial community metagenomes. By using QuantMeta, researchers can quantify a wider range of genes or populations simultaneously, as compared to conventional qPCR-based approaches.

**Figure 1.**
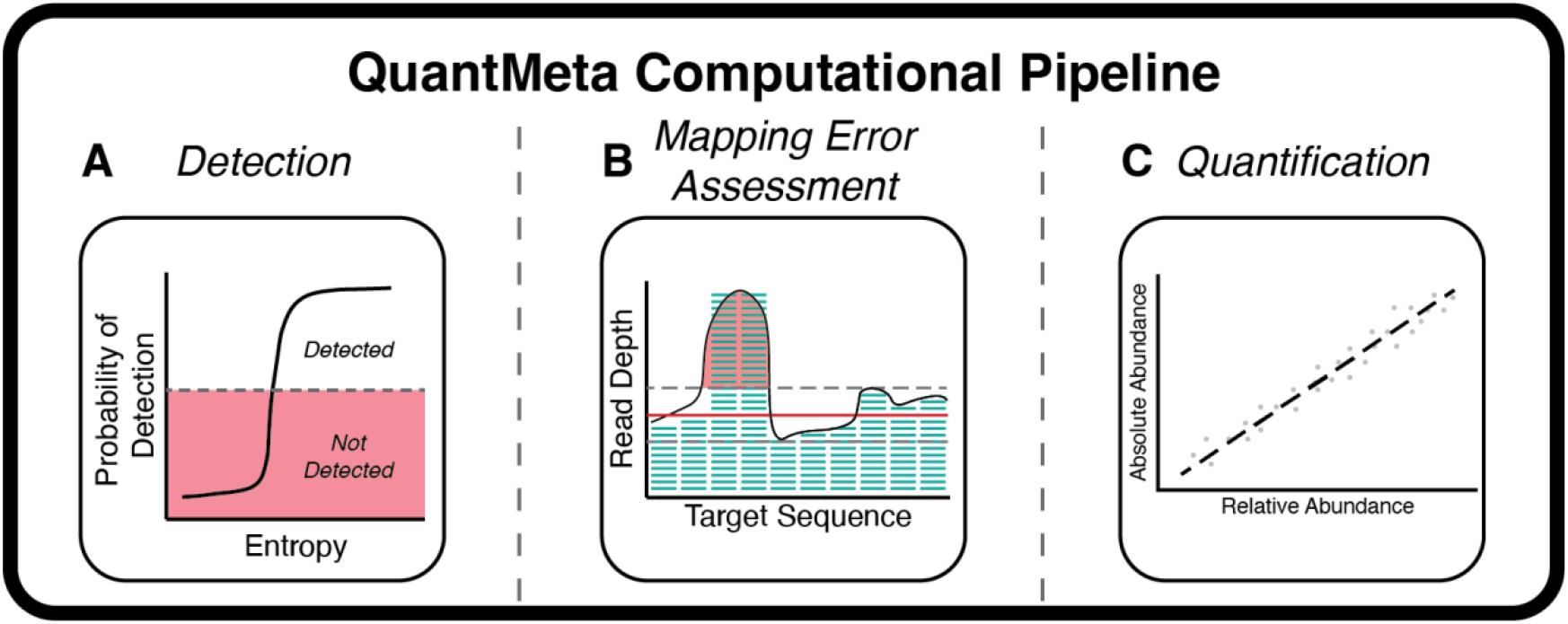
Conceptual overview describing the three main steps of the QuantMeta computational pipeline: detection, mapping error assessment, and quantification. (A) In detection, short reads mapped to standard sequences are used to establish detection thresholds. (B) In mapping error assessment, read depth variability thresholds are set for assessing read mapping errors. (C) A standard curve is created by relating relative to absolute abundances. Short reads mapped to target sequences from assemblies or databases are assessed for meeting the defined detection thresholds Read mapping errors are detected and corrected. The absolute abundances are then quantified.

## MATERIALS AND METHODS

Several terms used consistently throughout our work are defined in SI Section 2.

### Sample Collection and Processing

Two grab samples of secondary effluent (20 L) or raw influent (10 L) were collected in autoclaved carboys from automatic samplers at the Ann Arbor Wastewater Treatment Plant (Ann Arbor, MI, USA) daily from December 19 to 24, 2020. Samples were transported to the lab on ice within 2 h of collection. They immediately underwent concentration and purification steps based on a previously described ultrafiltration method (23). Briefly, secondary effluent and raw influent were pre-filtered through 100-μm sized pores (Long-Life polyester felt filter bag, McMaster-Carr, Cat. No. 6835K58) then 0.45-μm sized pores (Express PLUS PES filters, MilliporeSigma™, Cat. No. HPWP14250) and concentrated with tangential ultrafiltration (approximately 50-fold and 25-fold, respectively, using 30 kDa MWCO dialysis filters, Asahi Kosei Medical Co., Ltd, Cat. No. 6292966). The concentrated samples were treated with 1 mL chloroform to lyse any remaining cells, filtered through 0.45-μm filters, further concentrated with dead-end ultrafiltration (approximately 20-fold, using 100 kDa MWCO Amicon™ Ultra Centrifugal filter units, MilliporeSigma™, Cat. No. UFC510096). The extra-viral nucleic acids were then degraded with 100 U/mL DNase (Roche, Cat. No. 10104159001) for 1 h on the bench top. The DNase enzymatic reaction was ended by adding 100 mM EDTA and 100 mM EGTA. DNA was then extracted immediately with QIAamp UltraSens Virus Kit (QIAGEN, Cat. No. 53706). The manufacturer’s instructions were followed, except the first six steps were replaced by combining 140 μL of sample with 5.6 μL of carrier RNA and briefly vortexing.DNA extracts were stored at -20°C. On December 26, 2020, a 20 L deionized water sample was processed with the same viral enrichment process to account for contamination during sample processing.

### Sample Characterization

Each raw influent and secondary effluent sample was analyzed for pH, turbidity, solids content, fold concentration, and viral recovery (Table S1). Sample volumes were determined by weighing samples and assuming a density of 1 g/mL. To determine virus recovery through enrichment, a replicate grab sample was always collected and processed in parallel with the grab samples intended for sequencing. The sample-specific viral recovery was determined by adding *Enterobacteria* phage T3 (GenBank accession no. NC_003298, ATCC^®^ BAA-1025-B1™) at a concentration of 10^5^ copies/μL. The concentration of phage T3 remaining after the enrichment steps was determined using a ddPCR probe assay (23) (Table S2). The sample-specific viral recoveries ranged 22-45%.

### Sequencing Standards and ssDNA Standard Development

We used both Sequins dsDNA standards obtained from Hardwick et al. (2018) and ssDNA standards prepared in our laboratories. A total of 91 synthetic DNA standards were selected to capture the expected viral genome diversity, including GC content, sequence length, concentration, and DNA structure (double-stranded or single-stranded). Sequins metagenome mix A consisted of dsDNA standards with lengths varying 981 to 9,120 bp and GC content ranging from 20 to 71% (6). The dsDNA concentration of the Sequin metagenome mix A was measured with the Qubit™ dsDNA HS Assay (ThermoFisher Scientific, Cat. No. Q32851) with 2 μL of DNA template added to the 200 μL assay.

To capture DNA structural diversity, this set was expanded with our addition of five ssDNA standards. To design these, we applied a similar approach as was used for the sequin metagenome mix development. We inverted ssDNA viral genomes in the RefSeq virus database (downloaded on 3/19/2019) and fragmented the genomes into 1-kb long segments. The fragments were mapped to the NCBI nr database (downloaded on 8/15/2019) with Bowtie2 (v2.3.5). Inverted genome fragments with no alignments to the NCBI nr database were selected as candidate ssDNA standards. A random selection of five sequences from the potential candidates were selected with varying GC contents (31.7, 40, 45, 50, and 60%; Table S3) and made into Megamer^®^ ssDNA fragments without the complementary strand (IDT, Coralville, IA, Table S3). ssDNA standards were limited to five sequences and a length of 1,000 bp due to cost and synthesis constraints.

The ssDNA fragments were resuspended in molecular biology grade ddH_2_O (Fisher Scientific, Cat. No. BP28191) to an approximate concentration of 7.5 ng/μL, aliquoted into 10 μL increments, and stored at -20°C. Standards underwent a maximum of one freeze-thaw cycle. Immediately before use, ssDNA concentrations were measured with the Qubit™ ssDNA Assay (ThermoFisher Scientific, Cat. No. Q10212). A mix of the ssDNA standards was prepared to match the concentration ranges in the sequins metagenome mix A, from 10^4^ to 10^8^ copies/μL. The final concentrations of the ssDNA standards in the mix were 10^7^, 10^4^, 10^6^, 10^8^, 10^5^ copies/μL for the ssDNA standards with GC content of 31.7, 40, 45, 50, and 60%, respectively.

### Spike Addition of Standards and Foreign Marine Phage HM1 Genome

We spiked the standard mixtures and marine phage HM1 genomes into nucleic acid extracts from the three influent and three effluent samples. To assess the variability in quantification resulting from the spike-addition and sequencing steps, we created three technical replicates for one influent sample (12/21/2021) and one effluent sample (12/22/2021) spiking each technical replicates separately with the standards and HM1 genomes.

The sequins dsDNA metagenome mix A and the ssDNA standard mix were spiked into each sample nucleic acid extract to achieve 10 copies/ng sample extract DNA for the standards at the lowest abundance in the mixes. We predicted that the standards spiked at 10 copies/ng DNA extract would be near the detection threshold at our sequencing depth (approximately 200 million reads per sample) based on a previous observation that a single read was approximately 50 copies/ng DNA extract with a sequencing depth of 50 million reads (12). The spike-in absolute abundances were confirmed with ddPCR assays (details provided below).

To further examine quantification accuracy of low abundance viruses at our sequencing depth, we spiked in marine phage HM1 genomes (Genbank accession no. KF302034.1) into the sample extracts. This virus is foreign to our wastewater samples and the dsDNA HM1 genome (129,401 bps) has an average GC content of 35.7%. HM1 DNA was extracted using the QIAamp UltraSens Virus Kit with the modified protocol described above. Then, 15 μL of DNA extract with 1 μL of loading dye was run on a 0.3% agarose gel with 1 μL of 10,000x SYBR™ Gold nucleic acid gel stain (Invitrogen™, Cat. No. S11494) per 10 mL of gel at 3 V cm^-1^ for 90 minutes. GeneRuler High Range DNA ladder (Thermo Scientific™, Cat. No. FERSM1351) was run according to the manufacturer instructions. HM1 genomes were extracted from the gel with the QIAEX II Gel Extraction Kit (QIAGEN, Cat. No. 20021). The concentration of the purified HM1 genomes were measured with the Qubit dsDNA HS Assay. HM1 genomes were spiked into all sample nucleic acid extracts at an abundance of approximately 50 copies/ng DNA to reflect viruses in wastewater at lower concentrations. The HM1 genome absolute abundance in the nucleic acid extract was checked with ddPCR (details provided below).

### Spike-in ddPCR Assays

The absolute abundances of one dsDNA standard, one ssDNA standard, and the HM1 genome in the spiked sample nucleic acid extracts were checked with singlet ddPCR reactions performed with the QX200 AutoDG Droplet Digital PCR System (Bio-Rad Laboratories, Inc., Hercules, CA). For each plate, at least two ddH_2_O negative controls and two positive controls from spike-in stocks were included. Specific primers and probes were developed for HM1, dsDNA standard S1106_MG_020_A, and the ssDNA standard with 45% GC content (Table S2). The 22 μL reactions were prepared with 11 μL of 2x ddPCR™ Supermix for Probes (No dUTP; Bio-Rad Laboratories, Inc., Cat. No. 1863023), 0.4 μM of all probes and primers, and 3 μL of template. Droplets were generated using the automated droplet generation oil for probes (Bio-Rad Laboratories, Inc., Cat. No. 1864110) to a 20 μL volume, then PCR was performed on the C1000 Touch™ Thermal Cycler (Bio-Rad Laboratories, Inc., Hercules, CA) immediately after droplet generation. The ssDNA assays consisted of 40 cycles of denaturation for 30 seconds at 95°C, annealing for 1 minute at 56°C, and extension for 2 minutes at 72°C, then enzyme deactivation for 5 minutes at 4°C and 5 minutes at 95°C, and a final hold at 4°C. The same PCR reaction for dsDNA assays was performed with an initial denaturation step at 95°C for 10 minutes. Plates were run on the droplet reader within 1 hour of PCR completion. For each ddPCR reaction, thresholds were set using a previously defined method that uses kernel density to categorize droplets as positive, negative, or rain (24). We reran any sample that exhibited more than 2 droplet clusters, more than 2.5% of droplets classified as rain, or less than 30% compartmentalization. Inhibition was checked for the ssDNA and dsDNA assays by running 10 and 100-fold dilutions on one influent and one effluent sample and was not found to significantly alter the determined concentration; therefore, 10-fold dilutions were used. Due to the low abundance of the HM1 genomes, we could not test for inhibition of this assay with the samples and 1- or 2-fold dilutions were used. ddH_2_O negative controls infrequently resulted in positive droplets, and the corresponding concentrations were always much lower than the concentrations in the samples.

### Illumina NovaSeq Sequencing

Libraries were prepared with the Accel-NGS^®^ 1S Plus DNA Library Kit (Swift Biosciences, Cat. No. 10024) using 50 ng DNA measured with an Agilent TapeStation. Samples were sequenced on the Illumina NovaSeq 600 with five samples sequenced per paired-end 500 cycle SP flow cell yielding 251-bp long reads. Library preparations and sequencing were conducted by the Advanced Genomics Core at the University of Michigan. Quality control was performed by trimming Illumina adaptors and an additional 15 bp from the rightmost and leftmost of each read to remove adaptors from the Accel-NGS^®^ 1S Plus DNA Library Kit, and reads were decontaminated of PhiX174 with BBDuk (BBTools, v37.64). Bases with quality scores less than 10 were trimmed from reads, then trimmed reads with quality scores less than 10 or lengths less than 100 bp were removed with BBDuk (Table S4).

### Oxford Nanopore Sequencing

We conducted long read sequencing to improve viral assemblies because viral genomes commonly have repetitive regions, high mutation rates, and contain host genome fragments (25-28). DNA extracts from each influent and effluent sample (n=6) without added standards or phage HM1 genomes were cleaned prior to library preparations with the Zymo Genomic DNA Clean and Concentrate-10 kit (Cat. No. D4011, Zymo Research Corporation). Long read libraries were prepared with the Ligation Sequencing Kit (Cat. No. SQK-LSK109, Oxford Nanopore Technologies) and barcoded with the Native Barcoding Expansion 1-12 (Cat. No. EXP-NBD104, Oxford Nanopore Technologies). The six samples were sequenced on two flow cells (R9.4.1, Cat. No. FLO-MIN106, Oxford Nanopore Technologies) (Table S4). Basecalling was performed using Guppy (v4.2.3) and called reads were classified as either pass or fail depending on their mean quality score (≥ 7). Library preparations, sequencing, and basecalling were conducted by the Advanced Genomics Core at the University of Michigan.

### Assemblies and Viral Sorting

We used two assembly methods and pooled the results with the technical replicates processed separately. Hybrid co-assemblies were performed with long reads and short reads from each sample separately with metaSPAdes (v3.15.2) using kmer sizes of 21, 33, 55, 77, 89, and 127. Long read only assemblies were performed with Flye (v2.8.3) for each sample separately followed by several polishing steps including four rounds of Racon (v1.4.10), one round of medaka (v1.3.2), and short-read error correction with pilon (v1.24) (25). Following both assemblies, the contigs for each sample were pooled and contigs less than 1,000-bp were removed with length and count statistics provided in Table S7. The remaining contigs were assessed for likelihood of viral or proviral origin. Five viral detection methods were run on the contigs. VirSorter (v1.0.6) (29) with the virome flag, VirSorter2 (v2.2.2) (30), VIBRANT (v1.2.1) (31) with the virome flag, DeepVirFinder (v1.0) (32), Kaiju (v1.8.0) (33), and CheckV (v1.0.1) (34) end-to-end were run to assess likelihood of each contig being viral or proviral. Potential viral and proviral contigs were identified using the results of the five viral detection methods with previously established rules (35). The viral contigs were binned using vRhyme (v1.1.0) (36) with a minimum contig length of 3,000 bp with the resulting bins were referred to as viral populations. Of the contigs greater than 3,000 bp, 76.2-83.3% of contigs were classified as viral. The total viral population counts ranged from 3,950 to 5,770 in our viromes (Table S7).

### Downsampling and Creation of False Positive Dataset

To create reliable detection thresholds using a binary logistic regression model, we downsampled the reads to produce standards that spanned the method’s detection limits. Downsampling was performed by randomly sampling 1% and 20% of reads with seqtk (v1.3). To engineer a false positive dataset, we first created a set of mutated standard sequences by simulating single nucleotide polymorphisms with Mutation Simulator (37). This involved five sequential rounds of mutations, with a 0.1 probability for single nucleotide polymorphisms and a 0.02 probability for 5 basepair-length insertions, deletions, duplications, inversions, and translocations. To identify standards that would result in false positives, we selected mutated sequences with a coverage less than 10% or expected read distribution less than 0.3. These were used in the logistic regression model dataset to capture the variability in mapping to targets that should not be detected.

### Mapping

We performed both reference-based and contig-based quantification. Quality controlled short reads from deinterleaved fastq formatted files were mapped using Bowtie2 (v2.4.2) with the default parameters. The resulting Bowtie2 sam files were converted to a tabular format showing the number of reads mapping to each basepair of a target using samtools depth (v1.11).

For reference-based quantification, we mapped reads to viruses from multiple sources: the NCBI viral database, Virsorter curated database, marine phage HM1 genomes, crAss-like viruses genomes, and DNA virus pathogen genomes. Bowtie2 indexes were built with default parameters for each reference database: NCBI viral database (downloaded 9/6/2021), VirSorter curated database (downloaded 5/5/2021), dsDNA and ssDNA standard sequences, mutated standard sequences, HM1 sequence, and RefSeq crAss-like phage database (downloaded 10/4/2021). For the NCBI viral database, indexes were constructed using entire viral genomes and then divided into individual genes. For DNA virus pathogens, complete genomes were compiled from ViPR (herpesviruses, n = 1099; poxviruses, n = 395; downloaded 7/22/2022; (38)) and from RefSeq (adenoviruses or mastadenoviruses, n = 15; papillomaviruses, n = 64; polyomaviruses, n = 6; parvoviruses, n = 2; Torque Teno Viruses, n = 18; bocaviruses, n = 3; downloaded 7/22/2022). Genomes with at least 90% ANI similarity and at least 70% coverage were then clustered with CheckV (v1.0.1) (34), resulting in 118 virus pathogen clusters (Table S6). One virus per cluster was selected as a representative to build Bowtie2 indexes with default parameters.

For contig-based quantification, we mapped reads onto contigs from our generated viral populations, dsDNA and ssDNA standard sequences, marine phage HM1 genomes, and crAss-like virus genomes. Viral populations from each sample were indexed using the “large-index” parameter. Minimap2 (v2.17) was used to map contigs onto standard sequences, HM1 genomes, and crAss-like virus genomes from databases using default parameters. To assess the effectiveness of our read depth variability thresholds in detecting assembly errors (as described below), we used Blastn (v2.9.0) with default settings to compare the sequence similarity of contigs derived from standards against a custom database of standard sequences.

### Quantifying Targets within Metagenomes

Next, the relative abundances measured in the metagenomes were converted to absolute abundances. We applied an approach similar to Hardwick et al. (2018) to relate the observed measured relative abundance of the spike-in standards and their known absolute abundances (Table S8) with a log linear regression model (Figure 1C, Equation 1).

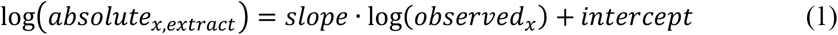

Here, absolute_x, extract_ (copies/μL) is the known absolute abundance in the DNA extract and observed_x_ (copies/μL) is the observed average read depths of a spike-in standard normalized to library inputs (Equation 2). Notably, observed_x_ of the ssDNA standards were doubled to account for having half as many sequences per copy of dsDNA.

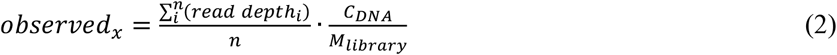

Here, read depth_i_ (copies) is the number of reads mapping to the i-th basepair along a target sequence with n basepairs, C_DNA_ (ng/μL) is the DNA concentration of the DNA extract, and M_library_ (ng) is the DNA mass used for library preparations.

A linear regression was created using the observed and absolute abundance of standards from all wastewater samples (Figure 2). The resulting regression was then applied to unknown targets in the same samples. The concentration of an unknown target in each sample (concentration_x,sample_, copies/mL) was then estimated by scaling the measured absolute abundance to the fold-change in volume that occurred during virus enrichment (Equation 3).

**Figure 2.**
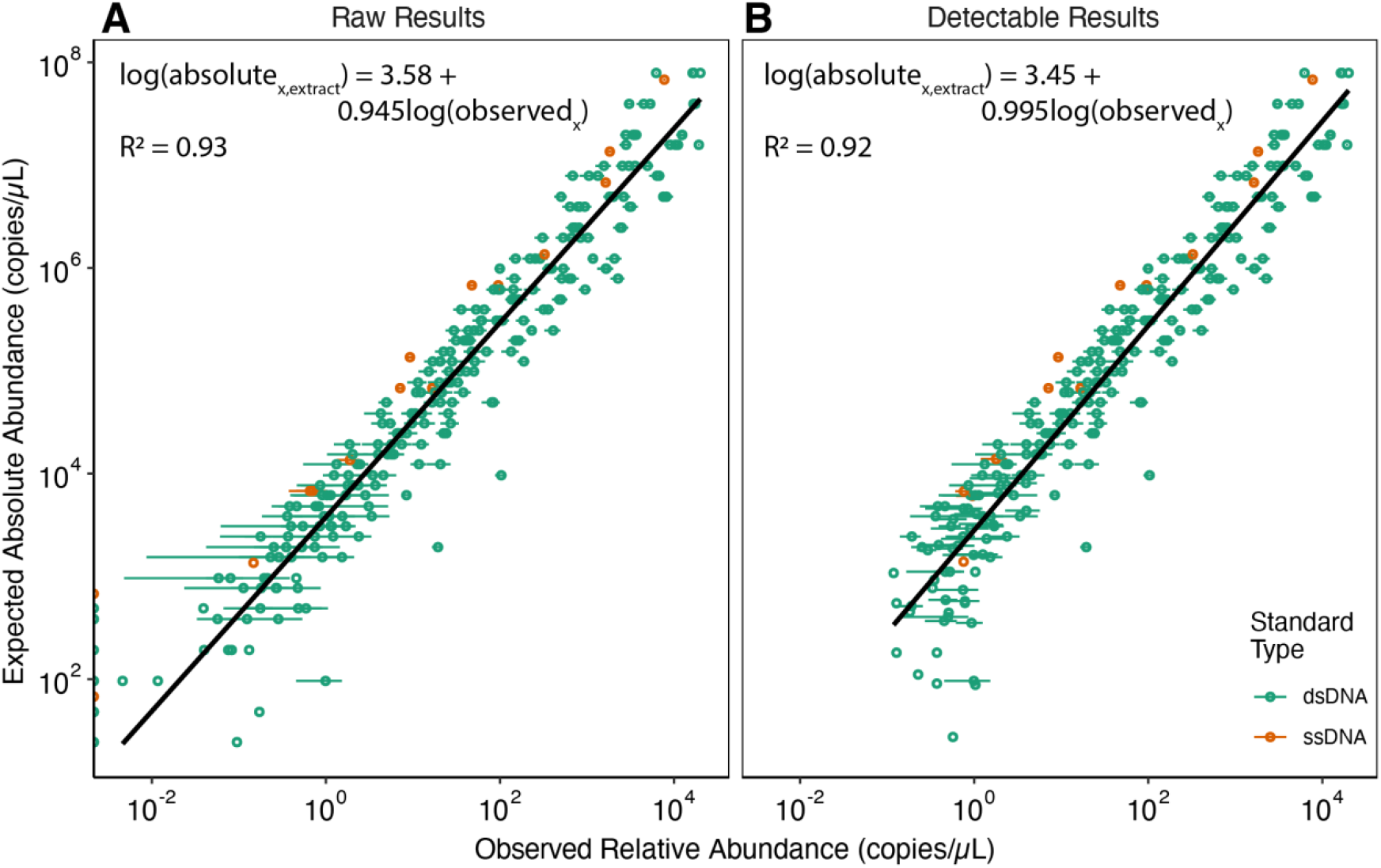
Application of the entropy-based detection threshold. The relationships between expected absolute abundances and observed relative abundances of the standards (A) before and (B) after the detection threshold was applied and standards that failed to meet the detection threshold were removed. The observed relative abundances per μL of DNA extract were determined with Equation 2 for all the standards across all samples, including the 20% and 1% downsampled results (mean relative abundance across all samples are reported as single values with standard deviations). The black line represents the linear regressions; green and orange points represent values of the dsDNA and ssDNA standards, respectively.

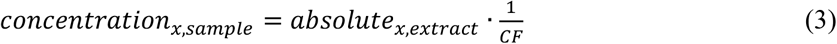

CF is the fold reduction of volume of the wastewater sample resulting from the viral enrichment steps. This approach assumes a 100% recovery of each target through the preanalytical virus enrichment and nucleic acid extraction steps. However, recoveries of our targets are likely less than 100%, as indicated by our phage T3 recovery controls. Currently, there is no comprehensive method to adjust target concentration of each target for preanalytical recoveries. Thus, it is standard practice to report concentrations assuming 100% recovery and provide measured recoveries for a control organism (39).

### Establishing Detection Thresholds

We next sought to establish thresholds for the detection of targets in a sample that considers both read distribution and coverage (as a proxy for abundance). We applied a metric, Shannon entropy, used as a diversity metric in ecology (40) that considers both species distributions and abundances. In our application, Shannon entropy is adopted as a strategy to relate read distribution and coverage in a single term (Equations 4-6, adapted from (41)).

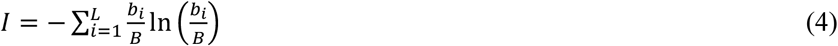

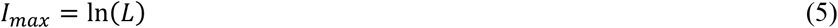

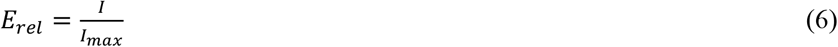

Here, I refers to the entropy of a target, while I_max_ represents the maximum possible entropy, indicating complete coverage and even read distribution for a target. The minimum value of I is 0, which represents when a single basepair from a read aligns to a target sequence. b_i_ is the read depth at basepair i along a target sequence. B is the total number of basepairs from reads mapping to the target. L is the length of a target sequence. Relative entropy, E_rel_, represents the evenness of read distribution, with 1 indicating complete coverage and perfectly even distribution of reads across the target sequence.

To assess how accurately relative entropy (E_rel_) reflects coverage and read distribution, we developed a binary logistic regression model. This model used reads mapped to both standard sequences and false positive sequences across all samples, including 20% and 1% downsampling. Detection was classified as pass or fail based on the cut-offs proposed by FastViromeExplorer (42), which require at least 10% coverage and an observed-to-expected read distribution ratio of 0.3 or higher. Unlike the FastViromeExplorer pipeline, we did not set a minimum allowable number of reads or basepairs from reads, due to the observed target length bias (Figure S3). The logistic regression model produced coefficients of ß_1_ = 320 and ß_0_ = -230 (Equation 7).

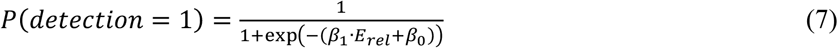

To determine the optimal entropy-based detection threshold, we first mapped reads to genomes in the VirSorter curated database, as well as genomes and genes from the NCBI DNA viral database. The combined mapping results from all databases yielded an area under the ROC curve of 0.992, indicating that the model has high sensitivity and specificity. The optimal entropy-based detection threshold (E_detect_) was determined using bootstrapping to maximize the combined values of sensitivity and specificity. The results suggested that the optimal E_detect_ threshold varied with target length. Targets were therefore binned by length and the optimal E_detect_ was calculated for each bin (Table S5).

### Statistical Analysis

All statistical analyses were performed in R (v4.0.3). The Wilkes-Shapiro test confirmed that neither relative nor absolute abundances followed a log-normal distribution (W > 0.95 and *p*-value < 0.01). Zero values were treated as NA, as the standards were designed to be evenly distributed across the concentration range used in the study. Linear and logistic regression analyses and student’s t-tests were performed with the R stats package (v4.0.3). Paired and unpaired t-tests were performed with 0.95 confidence levels with *p*-values less than 0.05 considered significant. ROC curves and optimal cutpoints for logistic regressions were calculated with the cutpointr package (v1.1.1). Shannon’s alpha diversity of viral populations in wastewater samples was determined based on viral concentrations in wastewater, with zero indicating non-detectable or absent populations. Graphs were created with ggplot2 (v3.3.5).

## RESULTS

We developed QuantMeta to determine absolute abundances of unknown targets from metagenomes with standard additions. To improve the accuracy of quantification, we established and assessed detection thresholds (Figure 1A), established read depth variability thresholds to detect and correct read mapping errors (Figure 1B), and then determined absolute abundances of viral targets in municipal wastewater samples (Figure 1C).

### Relationship between expected and observed abundances and spike-in reproducibility

Similar to earlier reports using synthetic DNA standards (6,8,11), we observed a strong linear relationship between observed relative abundances and expected absolute abundances of the standards (Figure 2A). The residuals of the ssDNA and dsDNA standards differed significantly (see SI Section 3 and Figure S2), likely due to differences incurred during library preparation or sequencing that are influenced by strandedness (43). However, the differences were small; when standard concentrations were predicted with either ssDNA or dsDNA standard curves, the absolute abundances differed between 5.9% and 12.5%. Therefore, we combined the ssDNA and dsDNA standards into a single standard curve for the remainder of the study.

We assessed the reproducibility of spiked standards in samples by sequencing technical replicates and conducting PERMANOVA tests. The relative abundances of each standard did not differ significantly among influent and effluent technical replicates (i.e., the standards added to replicate influent and effluent sample extracts; *p*-values = 0.18 and 0.36, respectively; Figure S1). In contrast, there were significant differences in the relative abundances of standards between different wastewater samples spiked with the same amount of standard among the three influent and three effluent samples (*p*-values = 0.0092 and 0.037, respectively). Moreover, the relative abundances of standards differed significantly between influent and effluent samples (*p*-value = 2.37 × 10^−9^). Across the six wastewater samples, the mean relative standard deviation for all standards was 21%, with higher relative standard deviations observed for standards spiked at lower abundances.

### One-metric Detection Threshold Improves Relative-to-Absolute Relationship

We observed poorer correlations between expected absolute abundances and observed relative abundances at the low range of standard concentrations (Figure 2A), suggesting the need for a detection threshold. Previous studies have set detection thresholds by setting a minimum read count (6,11,12) or incorporating read coverage and distributions thresholds (42,44). However, we found that implementing minimum read counts for detection thresholds resulted in target length biases, as a single read covers a larger proportion of shorter targets than longer targets (Figure S3). To address this, we developed a mapping entropy (i.e., measure of randomness) approach that incorporates both read distribution and coverage into a single detection threshold parameter (E_detect_, Figure 1A). This method is similar to how entropy is used to summarize community diversity and richness in the Shannon’s Diversity Index (45,46). We used standards and false positives to create a binary logistic regression model to generate probabilities of relative entropy (E_rel_) meeting read coverage and distribution requirements (Equation 7). We tested this with sample virome reads mapped to gene or whole genome targets from databases. A standard was confidently detected in the binary logistic regression if it had over 10% coverage and a ratio of expected Poisson read distribution to observed read distribution greater than 0.3; as empirically determined previously (42). Optimal entropy cutpoints were set to maximize the sum of sensitivity and specificity and varied with target length in our test dataset (Figure S4). Together, this resulted in a length-dependent entropy threshold, E_detect_ (Equation 8):

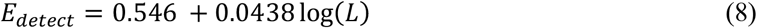

where E_detect_ is the unitless entropy threshold and L is the sequence length in basepairs. With this threshold, a target with a measured relative entropy (E_rel_) that is less than E_detect_ is not considered detected, as it does not achieve the minimum coverage and read distribution requirements.

The standards allowed us to identify detection limits in our wastewater viromes. Of 2,730 standards across all samples with 20%- and 1%-downsampling, 2,340 were detected and exceeded the length-specific detection threshold, resulting in a detection limit of approximately 500 copies/μL of DNA extract (Figure 2B). Standards that were not detected had a median expected absolute abundance of 96 copies/μL of DNA extract (range = 24-680 copies/μL). Removing these standards below the threshold substantially improved the relationship between the expected absolute abundance and observed relative abundances of standards. Ideally, the linear regression has a slope of one, indicating even sequencing across the range of absolute abundances. Without the threshold, the regression slope was 0.945 (R^2^ = 0.93); with the threshold applied, the slope improved to 0.995 (R^2^ = 0.92). We adopted the regression with the detection threshold to calculate absolute abundances of viruses in the wastewater extracts and estimate concentrations in the wastewater samples (see results below). The entropy detection thresholds in Equation 8 are applicable to other types of metagenomes, with or without standards added.

### Incorporating Read Mapping Correction Improves the Accuracy of Quantification

With a strategy to set detection thresholds in place, we used the standards to develop a method for detecting and correcting errors due to non-specific mapping and assembly errors (Figure 1B). When we quantified targets using reads mapped to *contigs* assembled from the sequenced standards, we observed poorer correlations between the expected absolute abundances and observed relative abundances (R^2^ = 0.67, Figure 3A) compared to mapping reads to known standard reference sequences (R^2^ = 0.92, Figure 2B). We attributed this discrepancy to assembly errors in the sequenced standards that interfered with standard read mapping (SI Section 5, Table S11).

**Figure 3.**
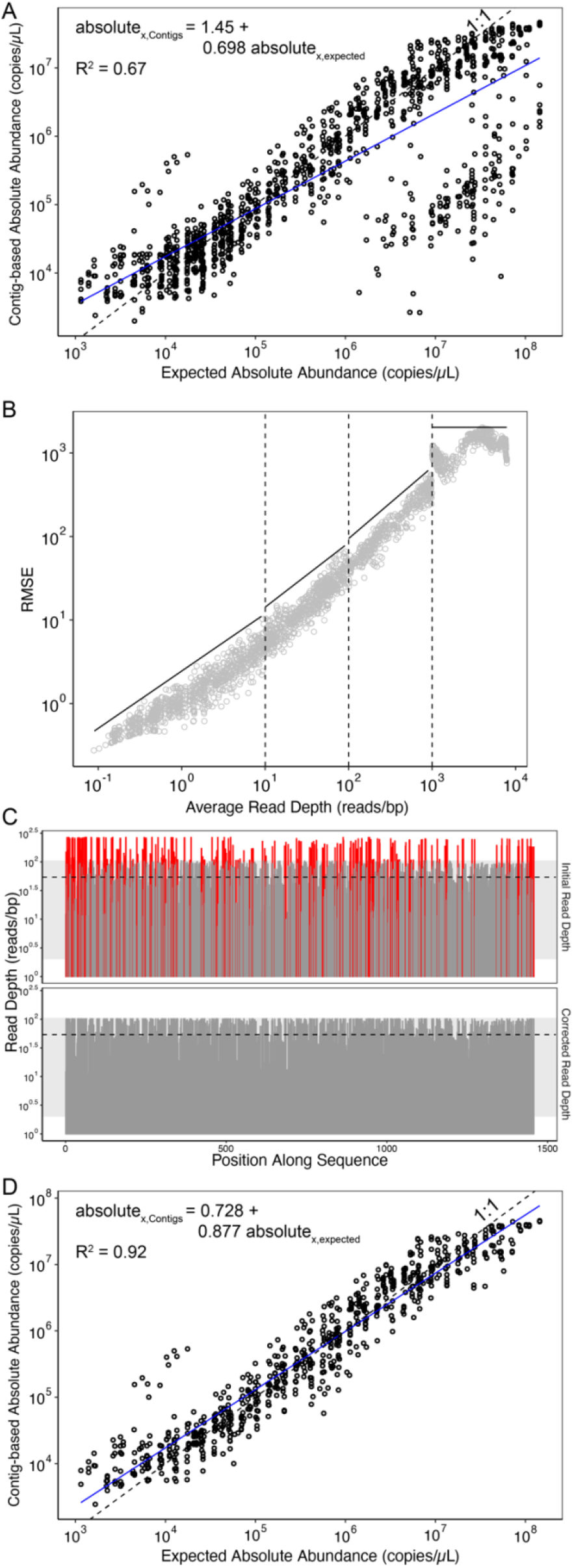
Quantitative accuracy improves by applying read mapping error assessment and correction. (A) Comparison of expected and contig-based absolute abundances of standards-derived contigs; each point represents a single standard-derived contig in a sample. The black, dashed line indicates the ideal 1:1 relationship between expected and contig-based quantification; the blue line indicates the observed linear regression. (B) RMSE thresholds (black lines) used to identify read mapping issues (e.g., non-specific mapping or assembly errors) based on RMSE of the reads mapped to standard references divided into bins at log_10_ intervals (dashed vertical lines, Table S3). (C) An example of a contig with nonspecific mapping or assembly errors before (top) and after (bottom) correction. Regions in red indicate areas outside 1.5 standard deviations from the mean read depth (gray shaded area). The corrected read depth is shown with the black, dashed line. (D) Comparison of expected and contig-based absolute abundances of standards-derived contigs after applying read mapping error assessment and correction.

In the absence of nonspecific mapping and assembly errors, the variability in the number of reads mapping across a target sequence is related to the local GC content and the read depth of the target sequence (47,48). Because the synthetic standards do not share DNA sequence homology with any known organism (6), we assumed there was no non-specific mapping to the standards. Thus, we used the relationship between local GC content and local read depth of the standard sequences (Table S9) to set maximum read depth variability thresholds for targets, where read depth variability is represented by root mean square error (RMSE) (Table S10, Figure 3B). Targets with read depth variability RMSE exceeding these maximum thresholds were identified as having either nonspecific mapping or assembly errors. They then passed through an iterative correction process (Figure S6): (1) identify outlier regions along the target (Figure 3C), (2) correct outlier regions (Figure 3C), (3) recalculate the target’s read depth variability RMSE (Figure 3C), and (4) iterate until the RMSE of the whole target is below the threshold or after 20 iterations. We stopped correcting after 20 iterations because most targets were corrected in less than 10 iterations, or did not achieve an RMSE below the threshold. If, after this process, targets had over 20% of 49-bp sliding windows altered or still had an RMSE above the threshold, they were deemed non-quantifiable. Further details on the design of these cut-offs are provided in the Supplementary Information (Figure S7).

To assess our ability to identify assembly errors with this approach, we evaluated the RMSE of reads mapping to the subset of contigs originating from the standards (Figure S8A). Of 910 standards in all samples, 827 were assembled into *de novo* contigs. Of these, 220 had read depth variability RMSE that was too high based on our thresholds and 21 of these were correctable. Correcting the standard contigs and removing the non-quantifiable standard contigs significantly reduced the differences between observed relative abundances and expected absolute abundances (*p*-value < 2.2×10^−16^). By evaluating the quality of the standard assemblies, we confirmed that the targets that exceeded our established read depth variability threshold indeed had assembly errors and that correction improved the quantification of those targets (SI Section 5).

### Quantification Errors Were Rare but Significantly Altered Target Concentrations

We applied the read mapping error assessment and correction approach to viruses that exceeded the detection threshold in the wastewater samples. Specifically, we identified read mapping errors amongst sequences from the NCBI RefSeq viral genome database (whole genomes and individual genes) and the VirSorter curated database (29), as well as representative sequences of viral populations assembled *de novo* from the viromes. In general, few targets exhibited read mapping errors. Specifically of the 162,139 genes and genomes from reference databases across all samples that exceeded detection thresholds, high read depth variability RMSE was observed for 3.6% of targets (n = 5,757) with 0.21% of targets (n = 338) identified as not quantifiable (Figure 4A). When the QuantMeta approach was performed on *de novo* assembled viral populations, the mean percentage of contigs with high read depth variability RMSE was higher than with reference databases, namely 11.1% (n = 14,849/133,322) across all samples (Figure 4B). On average across the samples, 4.2% of contigs underwent read mapping error correction with the remaining 6.9% of contigs remaining non-quantifiable.

**Figure 4.**
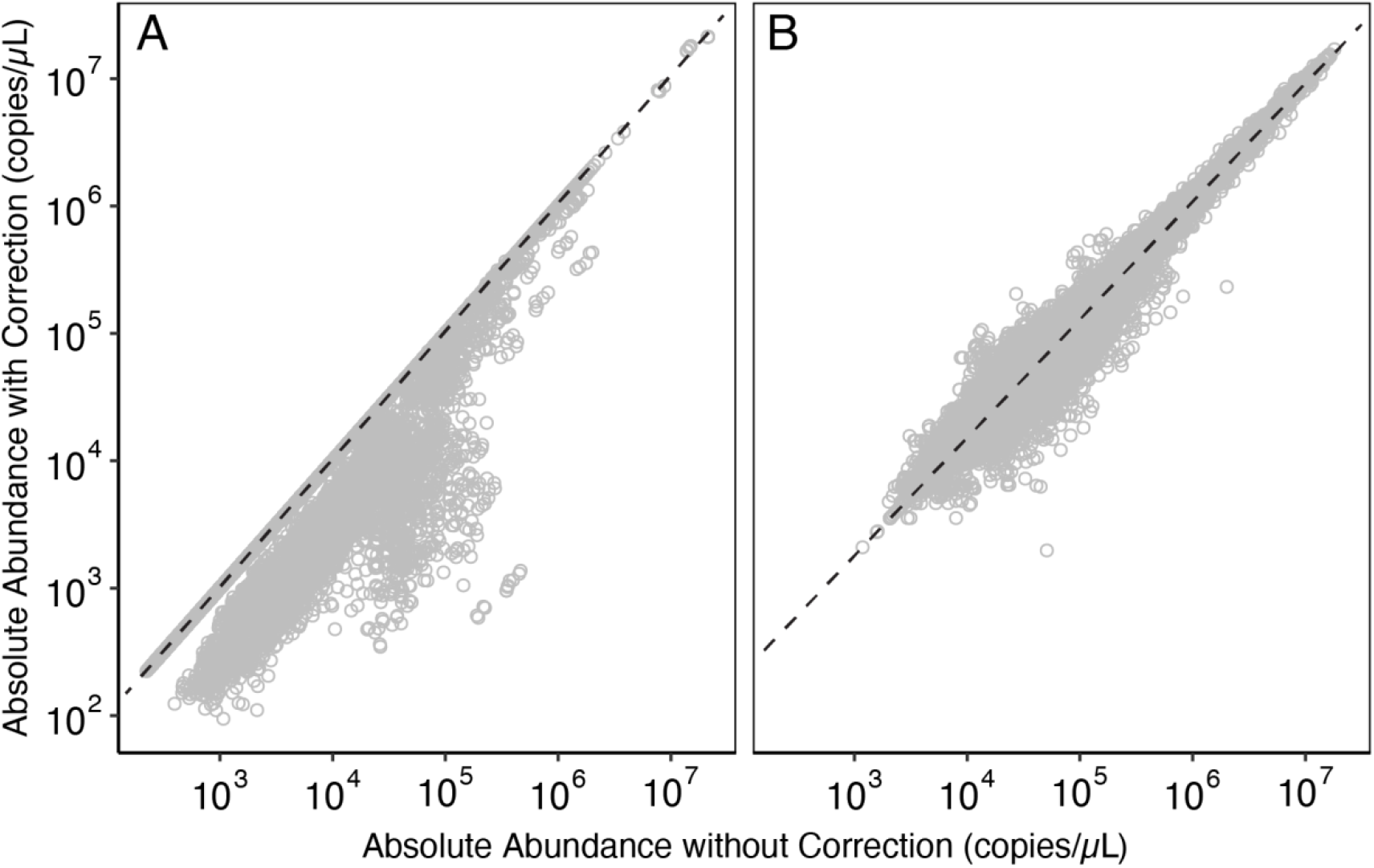
Impact of read mapping error correction on target absolute abundances. Relationship between target absolute abundances without (x-axis) and with (y-axis) read mapping error correction for reads mapped to viral sequences from reference databases (A) and *de novo* assembled viral populations (B). The databases included genomes and genes from the RefSeq viral and VirSorter curated databases. Targets that did not require read mapping error correction fall on the black, dashed line.

Targets deemed non-quantifiable after undergoing read mapping error correction are false positives within our metagenomes. The false positive rates (i.e., percent of reads mapped to non-quantifiable targets within reads mapped to each database) were 8.4%, 12%, and 22% for the RefSeq viral genes, RefSeq viral genomes, and curated VirSorter genomes, respectively. The false positive rate was 9.3% for viral contigs. These observations were consistent with previous read mapping to gene-centric databases that found false positive rates of 12-24% (49) and highlight the issues with quantification in the absence of read depth variability thresholds and correction.

Read mapping error correction impacted the observed absolute abundances of targets from database references (genes and genomes) and *de novo* assembled viral contigs differently. For the 3.6% of database targets undergoing correction, read mapping error correction significantly reduced the observed target absolute abundances (*p*-value < 2.2×10^−16^) by 65.2% (95% CI: 64.7 - 65.6%). The absolute abundances of *de novo* assembled viral populations were also significantly altered through the read mapping error correction process (*p*-value < 2.2×10^−16^), increasing or decreasing by 37.6% (95% CI: 37.5 - 37.8%) on average across the samples. For example, the concentration of *Salmonella enterica* phage LSPA1 (NCBI accession NC_026017, (50)) changed by an average of 2.8-fold following the correction process for the three samples flagged as having read mapping errors to the phage sequence. The targets that required read mapping error correction spanned all absolute abundances in our viromes.

### Viral Abundances Through Wastewater Treatment

#### Tracking Recovery of Spiked-in Phage HM1 Genome

To compare the QuantMeta approach with a standard molecular quantification method, we quantified spiked-in marine phage genomes (*Pseudoalteromonas* phage PSA-HM1) with both quantitative metagenomics and ddPCR. HM1 genomes were quantified by mapping reads to the HM1 reference genome (reference-based quantification) and by mapping reads to the *de novo* assembled HM1 contigs (contig-based quantification). The mean HM1 absolute abundance in the DNA extracts, as measured by ddPCR, was 1.1×10^3^ copies/μL (range: 7.2×10^2^-1.6×10^3^ copies/μL, Table S13). HM1 was consistently above detection thresholds and quantifiable by reference-based detection in all samples. Absolute abundances obtained via the QuantMeta reference-based approach closely matched those from ddPCR, differing by only 5.7% and 3.2% in influent and effluent samples, respectively. Technical replicates of HM1 reference-based absolute abundances among replicate influent and effluent samples were not statistically different (*p*-values = 0.011 and 0.019, respectively), suggesting that the QuantMeta approach is reproducible. *De novo* assembled HM1 contigs were identified in half of the samples, with the contig-based absolute abundances of HM1 generally exceeding those from ddPCR and reference-based measurements. Based on these HM1 contigs results and minimum absolute abundances of standards’ contigs (Figure 3D), a minimum absolute abundance of approximately 1,000 copies/μL is needed for successful assembly. Taken together, these results suggest that the detection limit of our contig-based quantification was approximately 1,000 copies/μL.

#### Total Virus Concentrations

We next applied the QuantMeta approach to quantify different groups of DNA virus populations commonly found in wastewater samples. Previous research has estimated total virus concentrations in wastewater based on epifluorescent microscopy and flow cytometry techniques. We measured total virus concentration by summing the concentrations of the identified *de novo* assembled and binned viral populations. Total virus concentrations were slightly higher in wastewater effluent than influent samples (*p*-value = 0.056), with mean concentrations of 1.5×10^9^ copies/mL (s.d. = 3.7×10^8^ copies/mL) in effluent and 7.5×10^8^ copies/mL (s.d. = 1.8×10^8^ copies/mL) in influent (Figure S9). The wastewater viral communities are diverse with mean Shannon alpha diversities of 7.9 (s.d. = 0.01) and 7.4 (s.d. = 0.05) in influent and effluent, respectively.

#### CrAss-like Phage Fecal Biomarker Quantification

CrAss-like phage, which infects Bacteroides in the human gut, is commonly measured in wastewater with PCR-based methods (51,52). We detected crAss-like phage in every sample, with higher concentrations in influent samples compared to effluent samples (Table 1). The mean reference-based measurements of crAss-like phage were 6.5×10^6^ copies/mL in influent (3/3 samples) and 1.8×10^5^ copies/mL in effluent (3/3 samples), with 1.7% of detected crAss-like phage sequences requiring read mapping error correction across all samples. The contig-based approach yielded higher concentrations, with crAss-like phage levels 154% higher than those from reference-based quantification (p-value = 0.067; Table 1). CrAss-like phage is commonly quantified in wastewater by qPCR using the CPQ056 primer set (53); however, this approach may underestimate crAss-like phage concentrations due to the specificity of the CPQ056 primers. When *in silico* quantification was limited to sequences targeted by the CPQ056 primers, crAss-like phage concentrations were reduced by 80-82% with reference-based quantification and 18-61% with contig-based quantification (Table 1).

**Table 1.** QuantMeta-derived concentrations of crAss-like phages (total crAss-like phages and CPQ056 primer-specific crAss-like phages) were measured in each sample by mapping contigs and reads to genomes from RefSeq. Concentrations are reported as copies/mL of wastewater. Concentrations did not account for viral recovery through sample processing; however, we observed 22-43% recovery of phage T3 through sample processing (Table S1).

| Sample | crAss-like phage/CPQ056 Concentration<br>(copies/mL wastewater) |  |
| --- | --- | --- |
|  | Contig-based | Reference-based |
| 12/19/20 Influent | $5.2 \times 10^7 / 2.0 \times 10^7$ | $7.1 \times 10^6 / 1.3 \times 10^6$ |
| 12/21/20 Influent | $5.0 \times 10^7 / 2.0 \times 10^7$ | $6.9 \times 10^6 / 1.3 \times 10^6$ |
| 12/23/20 Influent | $5.1 \times 10^7 / 2.1 \times 10^7$ | $5.6 \times 10^6 / 1.0 \times 10^6$ |
| 12/20/20 Effluent | $1.8 \times 10^6 / 1.5 \times 10^6$ | $2.3 \times 10^5 / 4.1 \times 10^4$ |
| 12/22/20 Effluent | $1.3 \times 10^6 / 1.0 \times 10^6$ | $1.8 \times 10^5 / 3.5 \times 10^4$ |
| 12/24/20 Effluent | $8.6 \times 10^5 / 6.0 \times 10^5$ | $1.2 \times 10^5 / 2.5 \times 10^4$ |

#### Human Pathogenic Viruses

Human pathogenic viruses are commonly measured in municipal wastewater with targeted PCR-based methods. We used QuantMeta to determine the concentrations of DNA virus pathogens in each of the wastewater samples (Figure 5). Whereas reads mapped onto the reference genomes of adenovirus, herpesvirus, bocavirus, papillomavirus, and polyomavirus, most of these targets in the samples did not meet our detection thresholds. Only polyomaviruses, papillomaviruses, and bocavirus met the detection thresholds and this was only in the influent samples except for one polyomavirus in one effluent sample. None of these required read mapping error correction (Table S13). For example, JC and BK polyomaviruses were quantified at 1.4×10^4^ and 1.4×10^4^ copies/mL, respectively, in influent samples. Most of the viruses failing detection were far from the detection threshold. Only 13 of 91 viruses below detection were within 10% of their respective detection threshold.

**Figure 5.**
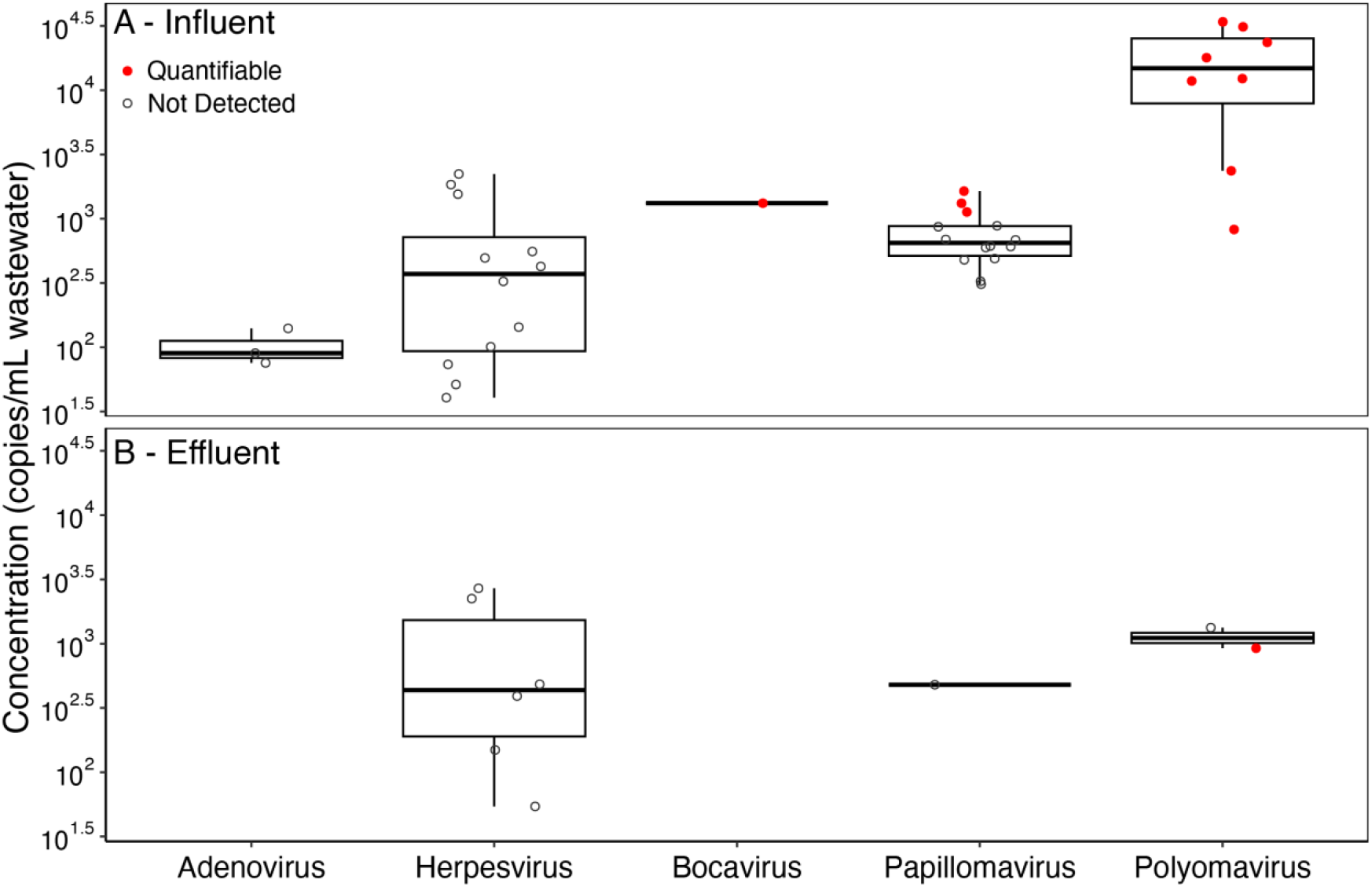
Reference-based concentrations of DNA virus pathogens were measured in influent (A) and effluent (B) samples by mapping reads onto representative genomes from clustering virus pathogen genomes in RefSeq and ViPR (see methods section for specifics). Each point represents the concentration of a virus cluster in a single sample with the status of passing or failing detection thresholds indicated in red and gray, respectively. Concentrations did not account for viral recovery through sample processing; however, we observed 22-43% recovery of phage T3 through sample processing (Table S1). All concentrations are provided in Table S13.

## DISCUSSION

Spike-in standards facilitate the quantification of microbial targets in metagenomes at accuracies similar to quantitative PCR methods (6-8,11,12). The major advantage of quantitative metagenomics is the ability to simultaneously quantify thousands of targets. Our research aims to enhance the reliability of these methods by rigorously defining their detection and quantification limits, thereby improving data trustworthiness. Specifically, we applied spike-in standards to establish detection thresholds and identify and correct read mapping errors in metagenomes (Figure 1). Incorporation of a new detection threshold and read mapping error correction improved the relationships between the expected absolute abundances and observed relative abundances of spike-in standards (Figures 2 and 3).

We assessed two methods of quantifying targets: (1) reference-based, by mapping reads to database reference sequences, and (2) contig-based, by mapping reads to *de novo* assembled contigs. To assess the detection threshold (i.e. minimum detectable absolute abundance) of these two approaches, we quantified the spike-in standards and a bacteriophage genome that was spiked into samples at low concentrations. For reference-based quantification, the entropy-based detection threshold (Equation 1) corresponded to an absolute abundance of approximately 500 copies/μL for the different targets. With contig-based quantification, the absolute abundance necessary to assemble a target was higher than that needed to exceed the entropy *detection* thresholds. Results from marine phage HM1 contigs and standards’ contigs suggest that the detection limit of our contig-based quantification was approximately 1,000 copies/μL. The minimum absolute abundances detectable with our approach will differ widely between studies and will depend on sequencing depth and sample complexity. Lower quantification thresholds are expected with lower community complexity and increased sequencing depth. Community complexity can be reduced with target enrichment steps, such as the virus enrichment performed on our wastewater samples to increase the virus-to-cellular genome ratio (23). The detection thresholds in our wastewater samples were approximately 100-fold lower than those previously reported in manure metagenomes (12), despite the previous study having less stringent thresholds (e.g., one read mapped required for detection). That study had four- to five-fold lower sequencing effort and more complex communities than our enriched wastewater viral communities. We note that our detection thresholds are higher than those of quantitative PCR methods, such as qPCR or ddPCR, which can be less than 10 copies/μL (54). Quantitative PCR therefore remains a more appropriate method for quantifying low abundance targets. As sequencing costs continue to decline, however, greater sequencing efforts and lower sequencing error rates will result in lower detection thresholds for metagenomics.

Our reference-based quantification involved mapping reads to database sequences representing previously sampled and sequenced population variants archived in a public sequence database. To quantify crAss-like phages using this approach, reads must closely match crAss-like phage genomes in RefSeq. In contrast, the contig-based quantification method maps reads to *de novo* contigs generated from population variants present in a sample. This method captures population variants that may be absent from databases, which could otherwise be missed in the reference-based method. Future applications of QuantMeta can incorporate varying levels of population variation when selecting references—such as contigs, metagenome-assembled genomes (MAGs), or sequences from databases like genomes or genes—to quantify targets. The choice of population variation depends on the specific goals of the study. For example, to quantify the concentration of a strain-level variant, reads would need to be mapped to a specific reference genome or to populations defined with minimal genome variation. Alternatively, quantifying a broader target group, such as all crAss-like phages, would involve using a reference population that accounts for greater population-level variation.

One of the most powerful applications of this approach on our samples was the ability to quantify total viruses in a sample. This has not been possible with quantitative metagenomic methods that rely on normalizing by viral counts or qPCR quantification. We addressed this challenge by incorporating single and double-stranded DNA spike-in synthetic standards in wastewater metagenomes that were highly purified for viruses. The measured total DNA virus concentrations in wastewater influent and effluent samples ranged 5×10^8^ to 2×10^9^ copies/mL, with effluent samples containing slightly more DNA viruses than influent samples. Previous studies using epifluorescent microscopy or flow cytometry to measure virus-like particle (VLP) concentrations in influent and effluent have reported similar or slightly lower concentrations, ranging from 1.0×10^8^-7.1×10^8^ VLP/mL and 1.0×10^8^-4.0×10^8^ VLP/mL in influent and effluent, respectively (55-57). These similarities in total virus concentrations across a variety of methods lends confidence to the reliability of the QuantMeta approach. The slight differences in total virus concentrations may be due to the reliance of epifluorescent microscopy and flow cytometry on intercalating dyes, which may underestimate counts of viruses with smaller genomes, including ssDNA viruses (58). Alternatively, the accuracy of the quantitative metagenomics approach depends on our ability to distinguish viral sequences from cellular genomes *in silico*. Therefore, the accuracy of this approach will continue to improve as our ability to identify viral sequences in metagenomic datasets improves (1).

The QuantMeta method allows for the simultaneous quantification of numerous viruses without introducing biases from PCR primers, which is particularly valuable for detecting human viruses in environmental samples such as municipal wastewater. In our study, we measured mean crAss-like phage concentrations of 6.5×10^6^ copies/mL in influent samples and 1.8×10^5^ copies/mL in effluent samples. By comparison, previous studies reported crAss-like phage marker gene concentrations ranging from 6.9×10^1^ to 1.1×10^9^ copies/mL in influent samples and 5.6×10^2^ to 1.0×10^6^ copies/mL in effluent samples (59-65). We hypothesized that the higher concentrations in our study were due to the limitations of the commonly used CPQ056 primer set, which was designed in 2017 based on just one of the now 129 available crAss-like phage genomes. This primer set may not capture the full genomic diversity of the crAss-like phage population in our samples. When we limited the *in silico* quantification of crAss-like phage to those detected by the CPQ056 primer set, the average crAss-like phage concentrations dropped by 81% for reference-based quantification and 42% for contig-based quantification (*p*-values = 0.012 and 0.072, respectively). This highlights a key advantage of quantitative metagenomics over qPCR or ddPCR: it avoids primer bias, which can miss portions of the target population. These findings are consistent with similar observations made when using quantitative metagenomics to quantify antibiotic resistance genes in agricultural systems (12).

When we measured DNA virus pathogen concentrations, reads mapped onto adenoviruses, herpesviruses, bocaviruses, papillomaviruses, and polyomaviruses (Figure 5). However, only a few pathogens achieved our detection thresholds: polyomaviruses, papillomaviruses, and bocavirus. Our quantitative metagenomics-based measurements of JC and BK polyomavirus concentrations (6.8×10^3^–2.1×10^4^ copies/mL) align with previously reported ranges for influent samples (8.9×10^0^–2.0×10^5^ copies/mL) (59-62,66). The wastewater concentration corresponding to our detection threshold (∼500 copies/μL of DNA extract) was relatively high compared to the concentrations of many pathogens in wastewater. Consequently, this quantitative metagenomics approach is not currently sensitive enough for wastewater-based epidemiology efforts that rely on sensitive detection thresholds (< 100 copies/mL). Additional sequencing effort would be necessary to confidently determine the presence and quantify other virus pathogens in the wastewater samples despite sequencing approximately 200 million reads per sample. The high detection thresholds of quantitative metagenomics relative to quantitative PCR methods impedes implementing the method to monitor pathogens in environmental matrices.

The QuantMeta method enables confident detection and precise quantification of populations in DNA extracts, offering significant utility for metagenomic studies. Our quantitative metagenomic approach is freely available as QuantMeta (github.com/klangenf/QuantMeta), with R notebooks that allow users to customize detection thresholds, read depth variability regressions, and thresholds to suit their specific applications. Users can apply more or less stringent entropy detection thresholds based on their own coverage and read distribution requirements. For example, Castro et al. (2018) provided parameters specifically for bacterial population detection, while KrakenUniq (67) offers an alternative k-mer based approach for reference-based detection. Technical variability can be further assessed with standards and replicates, as demonstrated in the DIVERS approach (68).

This approach has some limitations, particularly concerning sequencing depth, library preparation, and viral enrichment methods. For instance, the observed minimum absolute abundances, read depth variability regressions, and detection thresholds reported here may be specific to our sequencing setup—using Illumina NovaSeq SP flow cells with 251-bp paired-end reads and no amplification during library preparation. QuantMeta was designed for shotgun sequencing applications and is not suitable for amplicon sequencing. Our reported concentrations in the original wastewater samples are based on fold-changes through viral enrichment and nucleic acid extraction steps and do not account for the expected differential recovery of populations during processing (23). Prior research has addressed differential recovery by adding whole cells from various domains prior to sequencing (69). Although we included ssDNA standards, our linear regression model (Figure 2B) did not incorporate DNA structure, which may influence abundances. Despite these limitations, QuantMeta offers a robust and flexible tool for quantitative metagenomics, enabling more accurate assessments of microbial populations across diverse sample types.

## DATA AVAILABILITY

All sequencing data are available in the NCBI BioProject database under the accession PRJNA853368. All relevant code is compiled as QuantMeta and available on Github at github.com/klangenf/QuantMeta. Code generated for analyzing our wastewater viromes is available at github.com/klangenf/QuantMeta_Analysis.

## ACKNOWLEDGEMENTS

We thank Jacob Evans for improving the efficiency of the script listing the average GC content and read depths of 49-bp sliding windows and Eric Bastian for providing the viral RefSeq database divided into open reading frames. We thank the lead investigator of the NSF PIRE award, Peter Vikesland, for his leadership of the research study.

