## Supplementary material for "Development of a quantitative metagenomic approach to establish quantitative limits and its application to viruses": SI Figures

[1]Department of Civil and Environmental Engineering, University of Michigan
1351 Beal Ave, EWRE
Ann Arbor, MI 48109

[2]Department of Ecology and Evolutionary Biology, University of Michigan
1105 N University Ave
Ann Arbor, MI 48109

[+]Co-corresponding authors
Melissa Duhaime:
Krista Wigginton:

### SECTION 1: Summary Tables

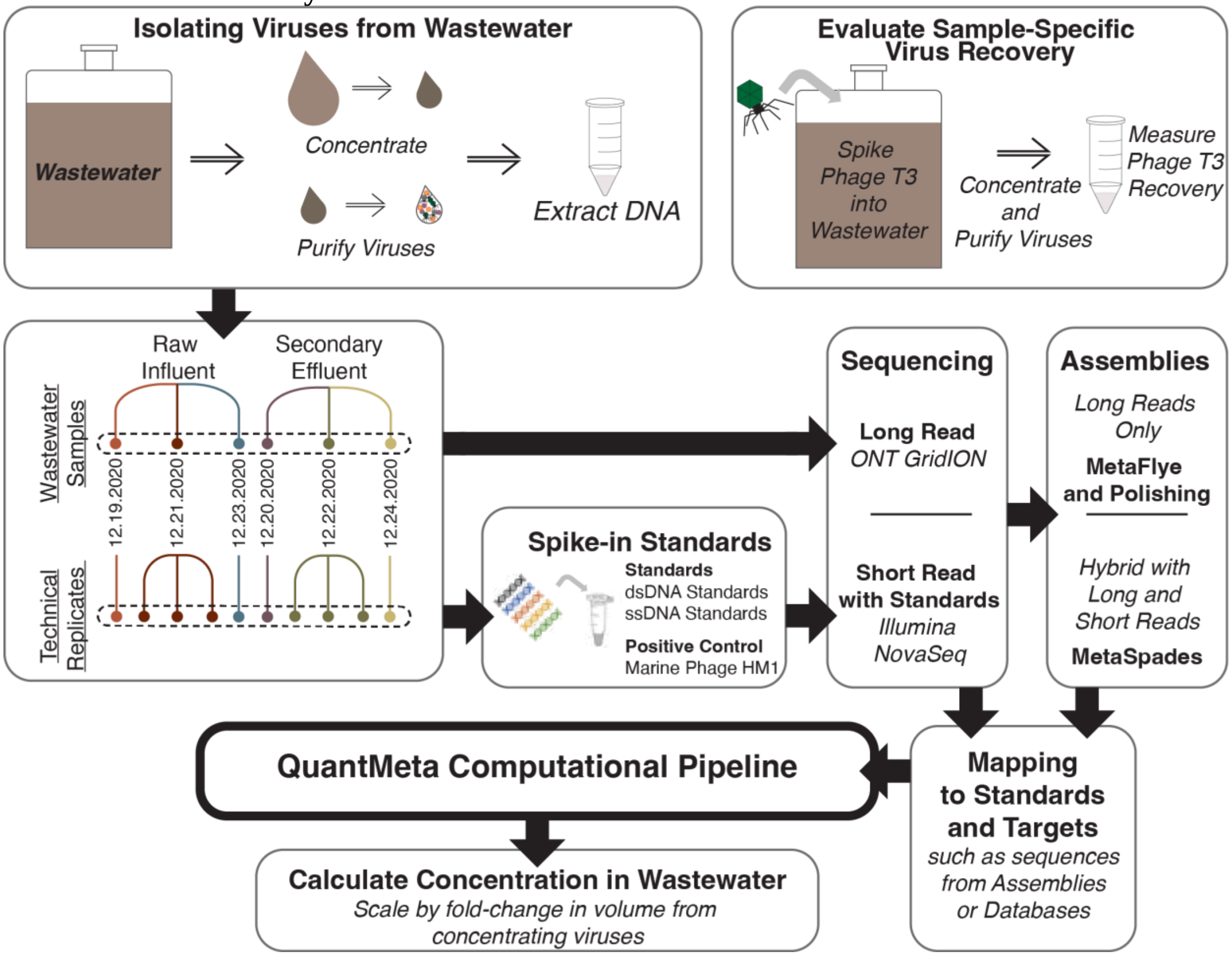

**Figure S1** Influent and effluent samples were concentrated and purified for DNA viruses. A separate sample processed in parallel was spiked with phage T3 to assess viral recovery. Viral DNA extracts were sequenced with technical replicates using long read and short read sequencing technologies. Synthetic DNA standards and marine phage HM1 genomes were spiked into samples before short read sequencing at known concentrations. Resulting short reads were mapped onto standard sequences to establish detection thresholds, set read depth variability thresholds for assessing read mapping errors, and create a standard curve by relating relative to absolute abundances. Unknown targets were evaluated for meeting detection thresholds and assessed for read mapping errors and then corrected. The absolute abundance was determined for unknown targets achieving detection thresholds without read mapping errors. Absolute abundance was converted to the concentration of targets in wastewater by scaling to the fold-volume change during virus isolation.

**Table S1** Sample characteristics. pH was measured with a Mettler Toledo pH meter calibrated immediately prior to measurement with 4, 7, and 10 pH standards. Turbidity was measured with a Hach 2100N Laboratory Turbidimeter. The total suspended solids (TSS) and volatile suspended solids (VSS) were measured in each sample using standard methods with 80 mL of sample stored at -20˚C until analysis (1).
**Table S2** ddPCR assays.
**Table S3** ssDNA standard sequences.
**Table S4** Spike-in concentrations and sequencing statistics.
**Table S5** Detection Threshold with Length Raw Data (for Figure S4).
**Table S6** Clusters of DNA virus pathogens. Pathogens were clustered by 90% ANI similarly and sharing 70% coverage using CheckV. A single virus genome was selected as a representative of the cluster.
**Table S7** Assembly and viral sorting statistics including contig lengths and number of contigs for hybrid and long read only assembly processes.
**Table S8** Individual Standard Concentrations.

**SECTION 2: Terminology**

Several terms used consistently throughout our work have definitions provided below.

Virome: Metagenome resulting from samples that were enriched for viruses.

Quantitative Metagenomics: A method to determine the absolute abundance of targets (e.g., genes, populations) in samples using metagenomics.

Relative Abundance: Quantity of a target relative to the other targets in a sequenced sample.

Absolute Abundance: Quantity of a target in a specific mass or volume of the sample (e.g., copies/µL DNA extract).

Concentration: Absolute abundance in the original sample (e.g., copies/mL wastewater) determined by converting absolute abundance in a DNA extract with changes in volume during sample processing.

Coverage: Fraction of a target sequence with reads mapped to it.

Read Depth: Number of reads mapped to a specific basepair along a target sequence.

Detection Threshold: Minimum entropy required for a target sequence to be confidently detected in a metagenome.

Read Depth Variability Threshold: Maximum read depth variability allowed across a target sequence for the absolute abundance to be determined accurately with quantitative metagenomics.

Reference-based Quantification: Determining the absolute abundance of target sequences in samples using sequences in databases.

Contig-based Quantification: Determining the absolute abundance of target sequences in samples using *de novo* assemblies.

**SECTION 3: Differences Between dsDNA and ssDNA Standard Regressions**

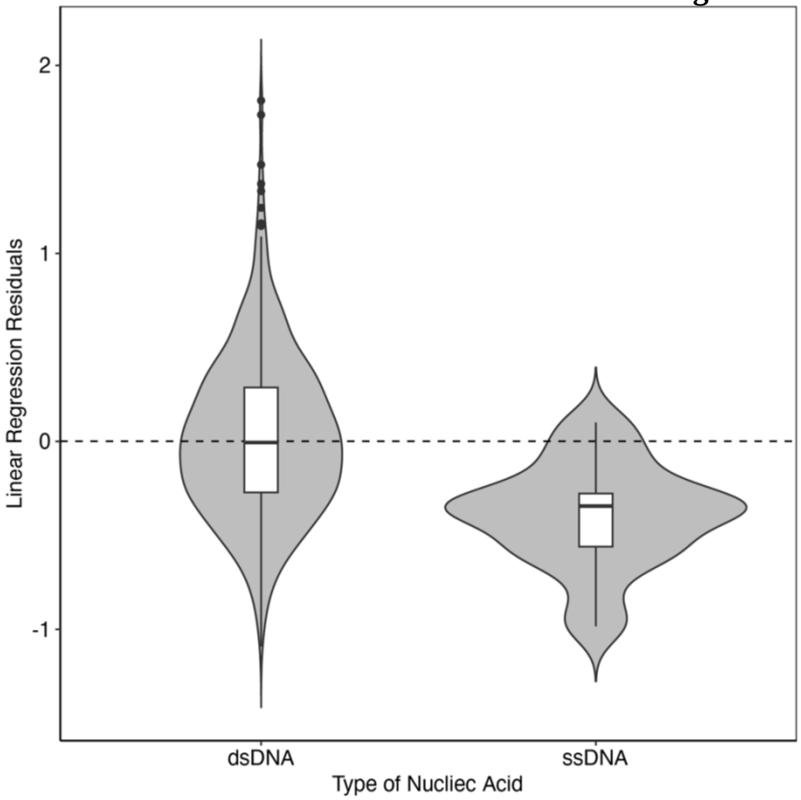


**Figure S2** The violin plot shows the different regression residuals of dsDNA and ssDNA standards across all samples to the linear regression relating expected absolute abundances to the predicted relative abundances. The residuals of dsDNA and ssDNA standards were significantly different (t-test, *p*-value = $5.6 \times 10^{-7}$). Furthermore, the additional consideration of DNA structure in an ANOVA significantly impacted the resulting linear regressions (*p*-value = $3.8 \times 10^{-7}$).

### SECTION 4: Establishing Detection Thresholds

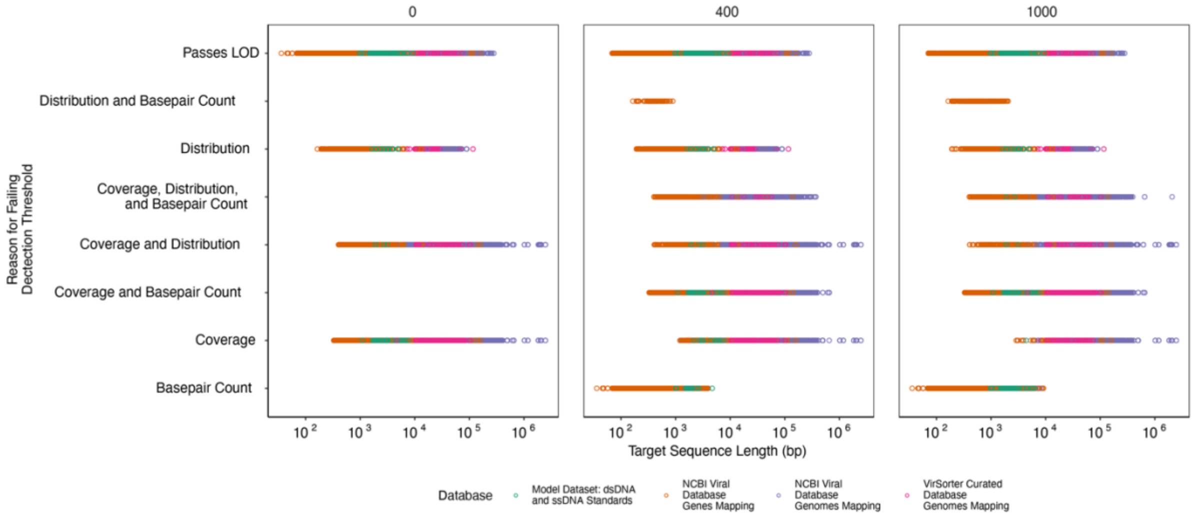


**Figure S3** Setting a minimum number of basepairs requirement introduced a sequence length bias to the detection of targets. Each panel provides reasons targets failed to meet detection thresholds when different minimum numbers of basepairs mapped to a target sequence were set (0, 400, and 1000 bp per target) with respect to the target sequence length. The results demonstrate that basepair count is redundant for long targets and only serves to increase the detection threshold of short targets.

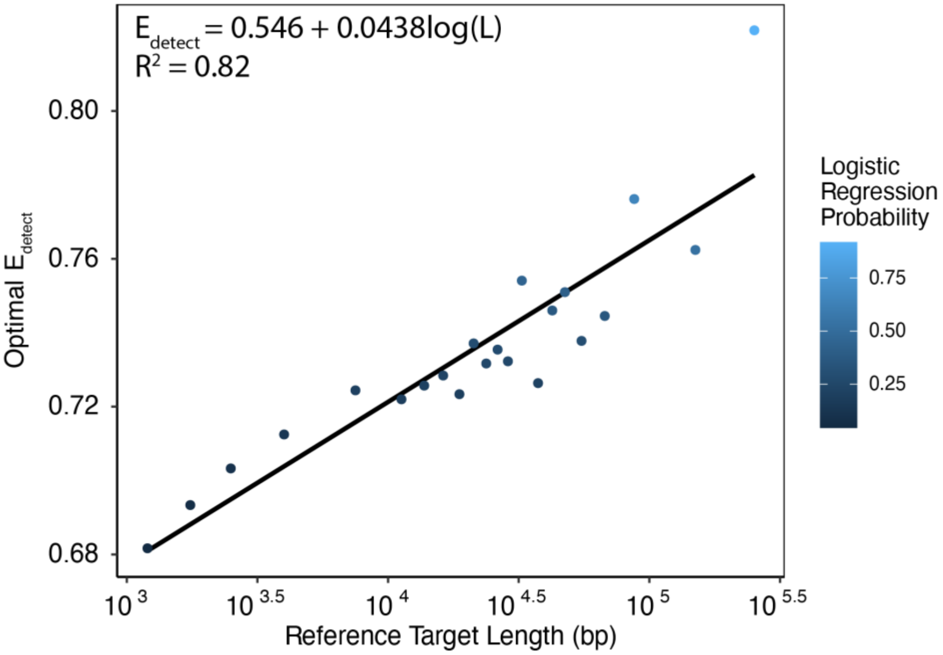


**Figure S4** Entropy-based detection threshold. The optimal $E_{detect}$ threshold was dependent on target length ($R^2$=0.82). Mapping results from all databases were pooled and divided into subsets based on sequence lengths with a minimum sequence length of 900 bp and a maximum length of 350,000 bp, then the optimal $E_{detect}$ was calculated for each length subset (Table S5).

### SECTION 5: Read Distribution Patterns Across Detected Targets

**Table S9** Read depth variability regressions. Regressions summarizing the variability in read depth along 49-bp long windows shifted by 1-bp for each spike-in standard with respect to the local GC content (i.e., average GC content of each 49-bp window). Separate regressions were developed for four ranges of average read depth with polynomial terms and $E_{rel}$ incorporated to prevent underfitting or overfitting. The normality of standardized residuals for each regression is plotted in Figure S5.

| Average Read Depth Range | Read Depth Variability Regression | $R^2$ |
|---|---|---|
| ≥1,000 reads/bp | $Read\ Depth_x = -22978 + 532(GC_x)^2 - 1745(GC_x) + 3426 \cdot \ln(Avg\ Read\ Depth)$ | 0.76 |
| 100 - 1,000 reads/bp | $\ln(Read\ Depth_x) = -0.090 + 0.371(GC_x)^2 - 0.185(GC_x) + 1.003 \cdot \ln(Avg\ Read\ Depth)$ | 0.83 |
| 10 - 100 reads/bp | $\ln(Read\ Depth_x + 1) = -0.676 - 0.154(GC_x)^2 + 0.230(GC_x) - 0.047 \cdot \ln(Avg\ Read\ Depth)^2 + 1.327 \cdot ln(Avg\ Read\ Depth)$ | 0.71 |
| 0 - 10 reads/bp | $\ln(Read\ Depth_x + 1) = 3.116 + 0.159(GC_x)^2 - 0.166(GC_x) + 0.134 \cdot \ln(Avg\ Read\ Depth)^2 + 0.351 \cdot \ln(Avg\ Read\ Depth) + 5.522(E_{rel})^2 - 7.799(E_{rel})$ | 0.57 |

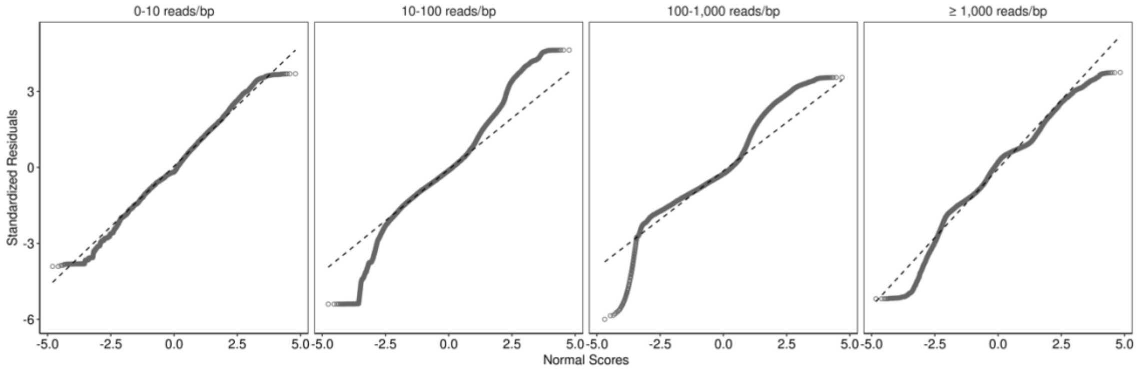


**Figure S5** Q-Q plots of read depth variability regressions. Each regression was amended to prevent overfitting and underfitting while striving for a normal distribution of standardized residuals. Q-Q plots were examined to assess the normal scores and that standardized residuals generally follow at 1:1 relationship.

**Table S10** Read depth variability thresholds. Acceptable read depth variability thresholds based on the root mean square error (RMSE) comparing the observed and predicted read depth variability for reads mapped to standard sequences. Separate thresholds were created for each read depth variability regression (Table S9). The thresholds are the linear trend lines through the standards with the highest RMSE translated up by $e^{0.25}$ (Figure 3B). Targets with a RMSE greater than the $RMSE_{max}$ at its average read depth are considered to have mapping errors.

| **Average Read Depth Range** | **Acceptable Read Depth Variability Threshold** |
|---|---|
| ≥1,000 reads/bp | $RMSE_{max} = 2027$ |
| 100 - 1,000 reads/bp | $\ln(RMSE_{max}) = 0.613 + 0.855(\ln(Avg\ Read\ Depth))$ |
| 10 - 100 reads/bp | $\ln(RMSE_{max}) = 0.880 + 0.771(\ln(Avg\ Read\ Depth))$ |
| 0 - 10 reads/bp | $\ln(RMSE_{max}) = 0.894 + 0.686(\ln(Avg\ Read\ Depth))$ |

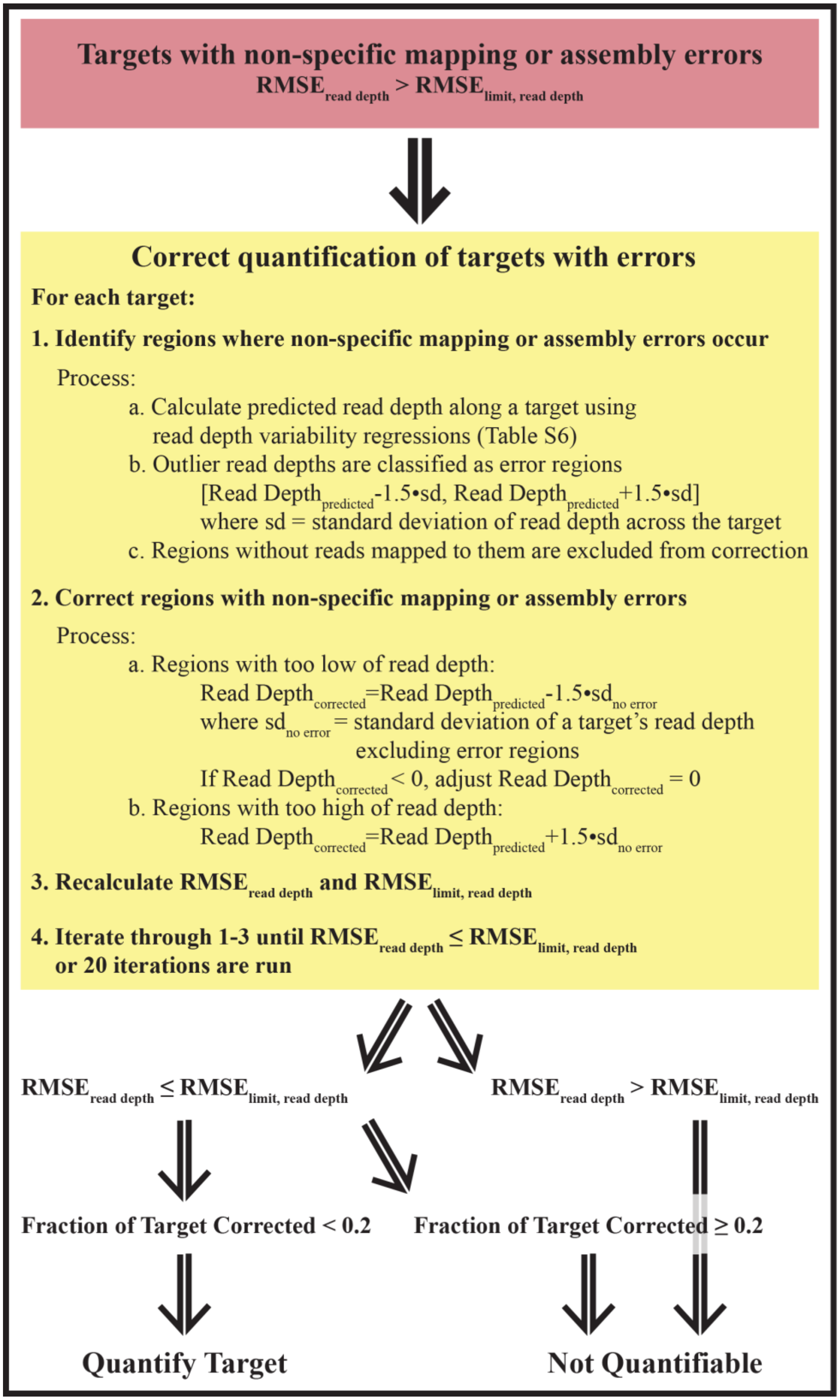


**Figure S6** Quantification correction approach. Overview of the method to detect and correct quantification of targets with non-specific mapping or assembly errors.

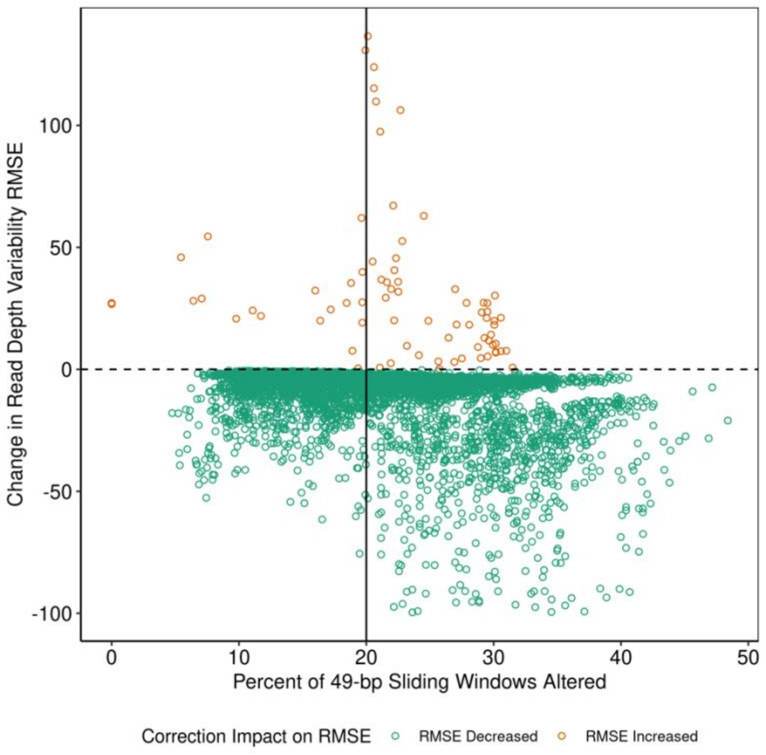


**Figure S7** Impact of quantification correction on RMSE. Observed changes in read depth variability RMSE with respect to the fraction of 49-bp sliding windows altered due to quantification correction. Increased RMSE (orange points) above 100 was observed for some targets when more than 20% of 49-bp sliding windows were corrected (black line). Targets with RMSE decreased more than -100 were excluded.

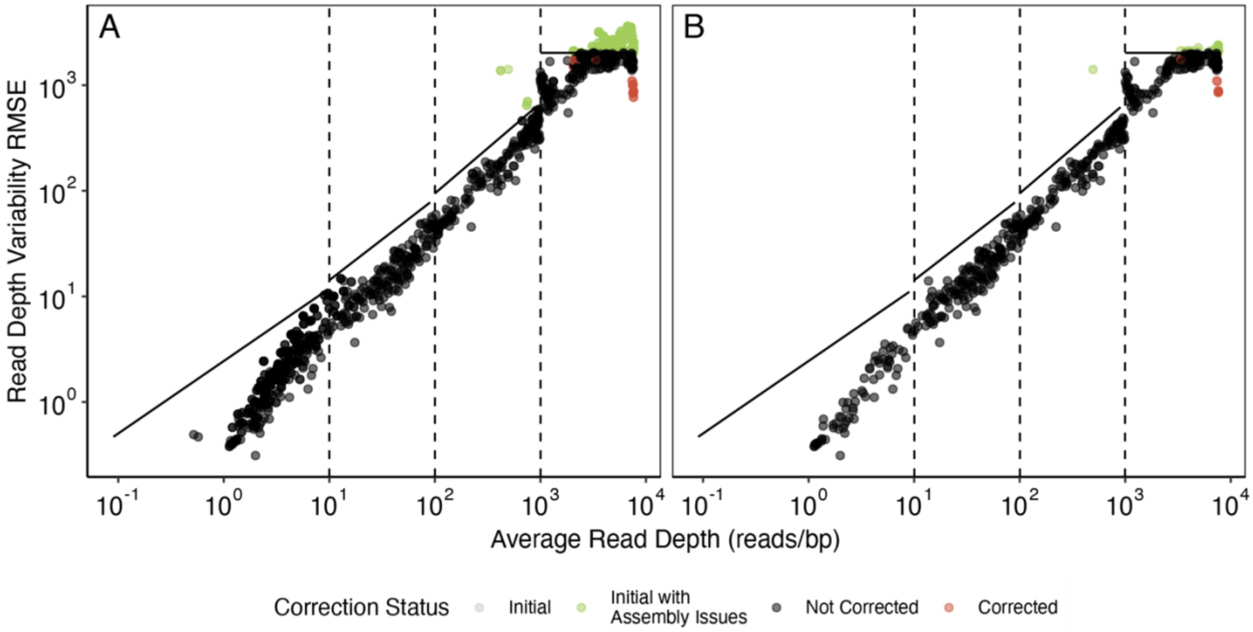


**Figure S8** Quantification correction detected assembly errors. Quantification error detection and correction applied to all contigs derived from spike-in standards (A) and coarse quality-controlled standards' contigs (B). Gray points represent initial RMSE with points falling above the black RMSE threshold lines requiring correction. Points in green are initial RMSE that were flagged as having assembly errors (fragmentation and low alignment). Corrected RMSE for all targets with less than 20% of sliding windows requiring correction are shown in red.

**Table S11** Contigs derived from standards with assembly errors were detected with the read depth variability threshold. Reads were mapped to standard-derived contigs with and without coarse quality control (i.e., redundant contigs of fragmented standard sequences or contigs with less than 80% alignment to the standard reference sequences). There were 26 standards' contigs with high read depth variability RMSE were not filtered by the coarse quality control measures. These standards' contigs had additions to the ends of the standard sequences on the contigs that impacted the read depth distribution. Incomplete and over-extended contigs were previously found to be unreliable estimates of transcript abundances (2).

| **Standard Derived Contig Statistic** | **All Standard Derived Contigs** | **Quality Controlled Standard Derived Contigs** |
|---|---|---|
| Number of Contigs | 1,234 | 568 |
| Number of Standards Represented | 827 | 568 |
| Number of Contigs with High Read Depth Variability RMSE | 220 | 26 (subset of the 220) |
| Number of Contigs Altered by Quality Control | 259 | 0 |
| Number of Contigs with High Read Depth Variability and Altered by Quality Control | 75 | 0 |

| | | |
|---|---|---|
| Number of Contigs with High Read Depth Variability and Not Altered by Quality Control | 26 | 26 (same 26 standards) |
| Number of Contigs without High Read Depth Variability and Altered by Quality Control | 184 | 0 |
| Number of Standards that were correctable | 21 | 8 |
| Number of Standards that were not quantifiable | 199 | 18 |

**SECTION 6: Reference-based and Contig-based Concentrations**

**Table S12** Marine phage HM1 concentrations. Measurements of marine phage HM1 genomes spiked in at a low concentration (~1000 copies/µL) into each DNA extract prior to Illumina NovaSeq sequencing. Contig-based virome derived measurements are not present in 5/10 samples (n.p. = not present).

| **Sample** | **Technical Replicate** | **Marine Phage HM1 Concentration (copies/µL DNA extract)** | | |
|---|---|---|---|---|
| | | **ddPCR** | **Contig-based** | **Reference-based** |
| 12/19/20 Influent | 1 | $9.17x10^2$ | $6.23x10^3$ | $1.53x10^3$ |
| | 1 | $8.62x10^2$ | n.p. | $9.08x10^2$ |
| 12/21/20 Influent | 2 | $8.76x10^2$ | $8.41x10^3$ | $1.19x10^3$ |
| | 3 | $1.11x10^3$ | n.p. | $1.33x10^3$ |
| 12/23/20 Influent | 1 | $7.23x10^2$ | $6.27x10^3$ | $1.76x10^3$ |
| 12/20/20 Effluent | 1 | $1.34x10^3$ | $9.91x10^3$ | $2.37x10^3$ |
| | 1 | $1.30x10^3$ | $9.05x10^3$ | $1.17x10^3$ |
| 12/22/20 Effluent | 2 | $1.28x10^3$ | n.p. | $1.33x10^3$ |
| | 3 | $1.07x10^3$ | n.p. | $1.83x10^3$ |
| 12/24/20 Effluent | 1 | $1.58x10^3$ | n.p. | $1.73x10^3$ |

**SECTION 7: Virome Quantification Results**

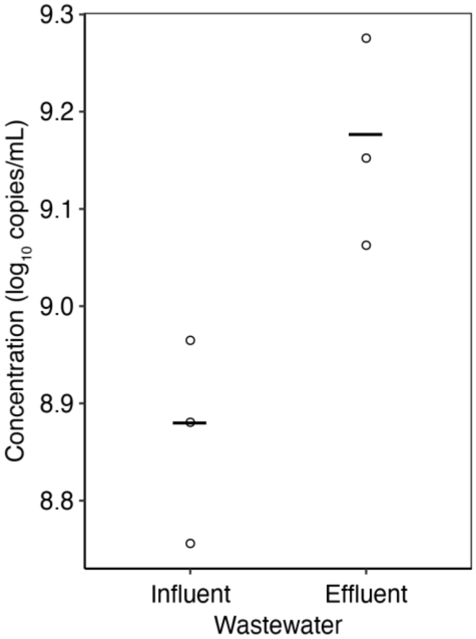


**Figure S9** Total virus concentrations. Concentrations of total viruses in influent and effluent samples as $\log_{10}$ copies/mL of wastewater. Dots represent viral population concentrations in individual samples with means indicated by black bars. Effluent does not have a significantly higher abundance of viruses than influent (*p*-value = 0.056). Concentrations did not account for viral recovery through sample processing; however, we observed 22-43% recovery of phage T3 through sample processing (Table S1).

**Table S13** Reference-based concentrations of virus pathogen clusters. Values in bold are quantifiable whereas all others were not above the detection thresholds (n.m. = no reads mapped to the target). Concentrations did not account for viral recovery through sample processing; however, we observed 22-43% recovery of phage T3 through sample processing (Table S1).
